# Rapid detection of *φx*-*174* virus based on synchronous fluorescence of Tryptophan

**DOI:** 10.1101/2021.11.27.470195

**Authors:** Yair Farber, Yaniv Shlosberg, Israel Schechter, Robert Armon

## Abstract

Development of rapid methods for detection of virus particles based on their intrinsic fluorescence is challenging. Pure viruses may be detected in filtered solutions, based on the strong fluorescence of the amino acid Tryptophan (Trp) in their proteins. Nevertheless, Trp also exists in high quantities in the hosts and host cultivation media. In this work, we show that a separation of the bacteriophage *φx*-*174* from its E. *coli* host (grown on the standard cultivation medium “Nutrient Agar”) by simple extraction and filtration is not sufficient for its detection based on the intrinsic fluorescence since ~70 % of the Trp fluorescence is derived from impurities. We formulate a new cultivation medium with very low Trp concentration. We apply synchronous fluorescence measurements to show that no Trp fluorescence is detected in the extract solution upon incubation of this medium substrate with ammonium acetate extraction buffer. Finally, we detect *φx*-*174* based on the spectral fingerprint of its intrinsic Trp content by synchronous fluorescence measurements. The concept of coupling intrinsic fluorescence-based methods to impurities reduction in the source, may pave the way towards future development of simple, cheap, and rapid methods for detection of viral pathogens.

## Introduction

Since the dawn of humanity, pathogens have always been considered as a major risk for the human health. Over the years, different efficient anti-bacterial agents were successfully developed such as anti-biotics and various peptide based drugs ^1^. However, in the case of viral attack, many infective diseases have remained cureless ^2^. In recent years, the global pandemic (caused by the SARS-CoV-2 virus) has spread worldwide, causing millions of deaths and severe illnesses ^3^. Rather than finding a cure, one of the main tools in the struggle against viral plagues is detecting the viral absence or presence in the population. Such a diagnosis enables quick isolation of the carrier, to prevent them from infecting additional people. Therefore, the development of sensitive and rapid viral detection methods continues to be of great importance.

Classical detection methods utilize bacterial or mammalian cells as hosts for the viruses. The cell lysis that is caused by the viruses forms discrete visible zones (plaques) that can be detected by eye or microscopic observation ^4^. Purification of viruses from their host cell debris and other contaminants is typically done by a variety of methods: density gradient centrifugation ^5^, ultrafiltration ^6^, or different chromatographic methods ^7,8^. The small sizes of viruses makes them invisible to simple microscopic observation and can be done by transmission electron microscopy (TEM) ^9^ or by utilization of fluorescent labeling agents under advanced microscopic instrumentation ^10^. Determination of viral structures was extensively investigated by X-ray crystallography, NMR and cryo-TEM ^11,12^. While a high purification degree of viruses is essential for their structural and fundamental studies, it may be time-consuming or require the utilization of expensive instruments. Therefore, such methods may not be feasible when a rapid detection is required. Identification of virus can be done by biosensors without an extensive purification process ^13–17^. In most cases, such biosensors are based on immunoassays in which unique antibodies can target specific viruses. The bound antibodies can be identified by simple gel-electrophoresis or immunofluorescence methods ^18^.

Another identification method that is very precise is the next generation sequencing (NGS) that can sequence the whole genome of the virus in a relatively short time ^19^. A big advantage of sequencing the entire viral genome derives from the ability to identify unexpected mutations and being able to follow evolutionary evolvement ^20^. Nevertheless, whole genome sequencing is still relatively expensive for large-scale applications.

A cheaper method for detection that is extensively used in recent years is polymerase chain reaction (PCR) ^11,12^. This method can detect viruses by amplification of specific sequences in the viral genome. An enhanced sensitivity was achieved by the development of loop-mediated isothermal amplification (LAMP) and recombinase polymerase amplification (RPA) ^21–25^.

Different Spectroscopic methods were reported to identify isolated viruses. Raman spectroscopy was reported to be able to identify viruses ^26^ and also help in determining and characterizing their structure ^27–29^. The amino acids Tryptophan, Tyrosine and Phenylalanine have an intrinsic fluorescence and can be used to detect isolated viruses ^30^. Intrinsic fluorescence can also be measured from nucleic acids, however, the intensity is relatively low, and in many cases, may not be sufficient to be detected ^31^. Nevertheless, the fluorescence intensity of nucleic acids can be detected by utilization of labeling agents such as SYBR dyes, that can significantly amplify it ^32^.

Several scanning methods are commonly available in commercial fluorimeters. The scanning method which is most extensively used is the emission scan in which the excitation is performed at a single wavelength and a wide range of emission wavelengths is collected. A more advanced scanning method is synchronous fluorescence (SF) ^33,34^. In this scanning method, both the excitation and emission wavelengths are scanned while keeping a constant wavelength interval (Δλ) between them. This is equivalent to scanning along a line of slope 1, in the 2D excitation-emission map. When the wavelength interval Δλ between the excitation and emission wavelength is chosen properly, the resulting spectrum will show one or several features that are much more resolvable than those in the conventional fluorescence emission scan. The maximum fluorescence intensity for a particular component occurs when Δλ corresponds to the difference between the wavelengths of the excitation and emission maxima for that component ^35,36^. SF spectra are often narrower than emission spectra and when mixtures are analyzed, selecting a proper Δλ may reveal more peaks and overlapping of multiple spectral components can be minimized ^37,38^. While SF methods have demonstrated as rapid tool for bacterial classification and identification ^39–42^, and even distinguishing between live and dead bacteria ^43^, to our best knowledge, no rapid detection of viruses by intrinsic SF has been reported.

Although fluorescence spectroscopy has a great potential for rapid detection of pure viruses based on the intrinsic fluorescence of their amino acids, it is not compatible for matrixes with many biological compounds that also consist of the same amino acids. In this work, we report a novel composition of a medium for the cultivation of the bacterial host of the virus *φx-174*, whose fluorescence contribution does not have any significant overlap with Trp. Also, we show that this medium enables the detection of the spectral fingerprint of *φx*-*174* by a simple filtration followed by SF.

## Materials and methods

### Replication of the virus *φx*-*174*

To replicate the virus *φx*-*174*, we used the “double-layer” method [44]. Briefly, in this method, a host bacterium (in high concentration) is grown into a solid medium together with infectious phage particles. After the phage is replicate itself in the bacterium, and lyse the bacterium cells, a visible clear zone (plaque) is formed. In our case, to E. *coli CN13* (ATCC 700609, a nalidixic acid-resistant strain) a *φx*-*174* (DSM 4497, Germany) was added and were grown on Nutrient Agar (NA) composed from Nutrient Broth (Difco, USA) with 0.7% Agar (Difco, USA). After overnight incubation (37°C) viral plaques were formed.

### Purification of the virus *φx*-*174* from whole E. *coli* cells

For separation of viral particles from whole E. *coli* cells, a 5 ml of Ammonium acetate buffer (Ammonium acetate, Riedel-de Haen, Germany, pH=7, 0.1M) were added on the “double-layer” Agar (Petri dish of 100 mm diameter). The buffer was float on the Agar for 10 min and then it pumped out by syringe and filtered by PVDF sterile syringe filter (0.22 μm, Millipore, Ireland). To estimate the contribution of lysed host bacterium cells and diffused molecules from NA on the SF measurements, the 0.22 μm filtrate were filtered again by 0.02 μm Alumina membrane (Anodisc filter, Whatman, Germany) and vacuum pump. The Alumina membrane were kept in dark and 4°C (for one overnight only) before tested for SF measurements in the day after.

### Preparation of LTM

The LTM consisted of Glucose, 10 gr/L (Merck, Germany); KH_2_PO_4_, 5 gr/L (Carlo Erba, Italy); K_2_HPO_4_.3H_2_O, 5 gr/L (Merck, Germany); (NH_4_)_2_HPO_4_, 3 gr/L (Merck, Germany); Na_2_HPO_4_, 4.6 gr/L (Fluka, Switzerland); MgSO_4_.7H_2_O, 0.6 gr/L, (Merck, Germany); Yeast extract, 0.5 gr/L (Difco, USA); Agar, 7 gr/L (Difco, USA).

### SF measurements

All SF measurements were performed using a Fluorolog 3 fluorimeter (Horiba) with excitation and emission slits bands of 7 nm at Δλ = 60 nm.

### Theoretical calculation of Trp units in *φx*-*174*

The theoretical number of Trp units in *φx*-*174* was calculated based on the sequences of the main proteins in *φx-17*: capsid protein F, major spike protein G, minor spike protein H, and DNA-binding protein J, that are deposited in the protein data bank (PDB [www.rcsb.org/pdb]).

### Preparation of Figures

Fig. 4 was created with BioRender.com

**Fig. 4.**
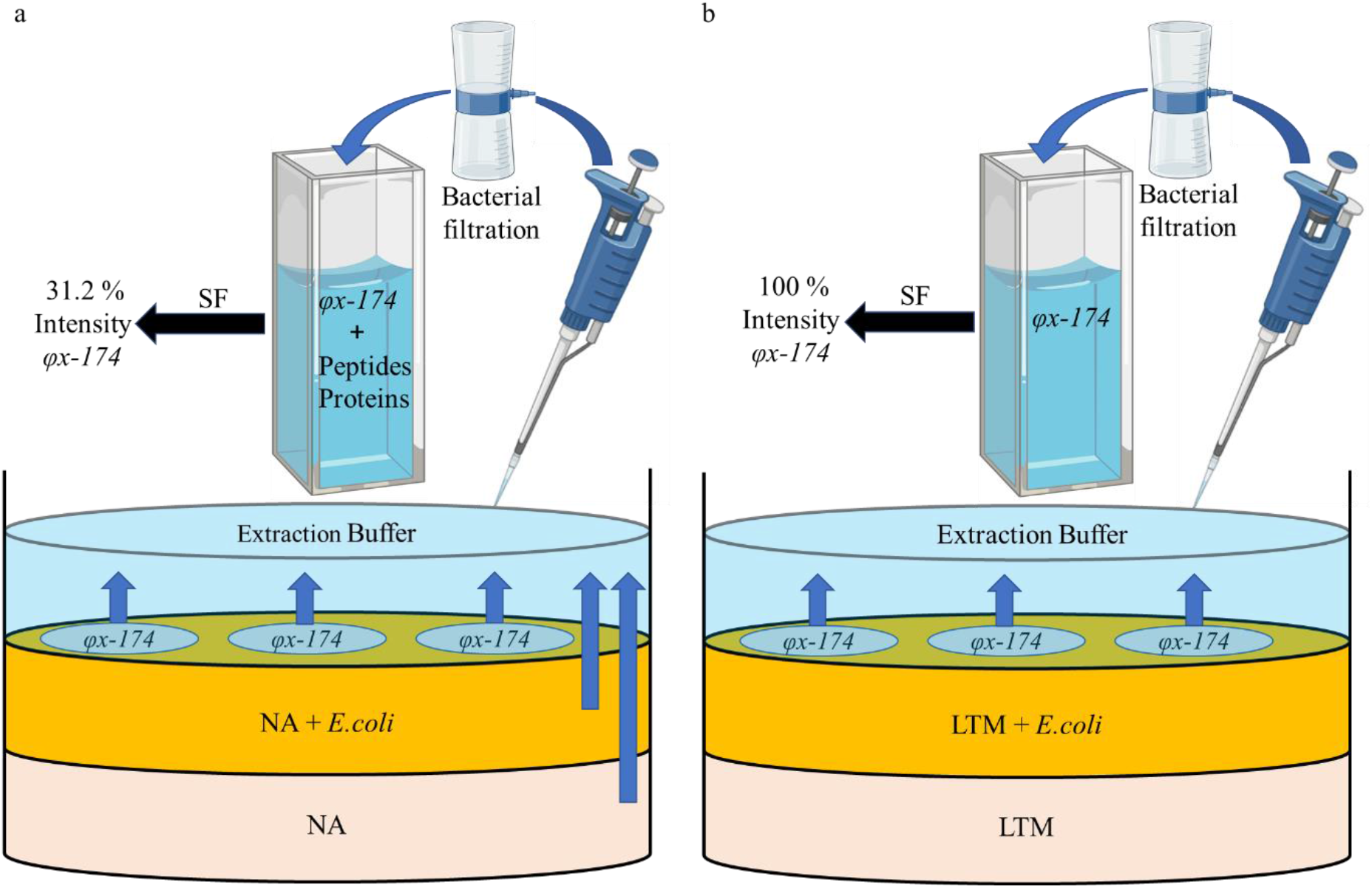
A model for the mechanism of the *φx*-*174* purification and detection. A schematic description of the method for purification and detection of *φx*-*174* that may work efficiently when the cultivation medium is LTM ^44^, but not on NA. *φx*-*174* plaques were replicated by the double layer method, In this method the bottom layer of the substrate is consisted of a bacterial cultivation medium (off white stack). The top layer is consisted of a bacterial host, mixed with bacterial cultivation medium (orange stack). The *φx*-*174* virus are replicated on top, forming plaques (blue circles). The virus is extracted by pouring ammonium acetate extraction buffer on top (cyan stack), followed by an incubation of 10 min. The extraction buffer is filtrated through a 0.22 μm filter (filter illustration) to remove whole bacterial traces. The detection of *φx*-*174* is conducted by SF measurements based on its intrinsic fluorescence of the amino acids Trp. The blue arrows represent the diffusion of components that consist of Trp towards the extraction buffer. A round arrow represents the filtration of the extraction buffer. Black arrows represent the SF measurements. The labels “% intensity *φx-174”* represent the % of the estimated Trp emission that derive from *φx*-*174*, in respect to the total emission at λ_em_ = 310 −370 nm. **a** The lower and top substrates are consisted of NA. The substrates enable the diffusion of proteins into the extraction buffer. Their spectra severely overlaps with *φx*-*174*, whose Trp emission intensity is estimated to be only 31.2 %. **b** The lower and top substrates are consisted of LTM. In this case, proteins and peptides are not extracted from the substrate. Therefore, 100 % of the Trp measured fluorescence derives from *φx*-*174* and may apply as a fingerprint for its detection.

## Results and Discussion

### Tryptophan residues in the cultivation media limit the detection of *φx*-*174* by Synchronous fluorescence

A frequently used method for replication of *φx*-*174* in Petri dishes is called the double agar layer method ^44^. In this method, the virus replicates within E. *coli* that is cultivated on top of layer of growth medium such as Nutrient Agar (NA). Whole bacterial cells can be easily separated from *φx*-*174* by simple filtration or centrifugation. However, peptides or proteins, which contain Trp, may also originate from lysed bacterial cells or diffuse from the lower layer of the cultivation media, making the purification *φx*-*174* more challenging. In our previous work, ^45^ we showed that the concentration of Trp can be directly determined by applying a single SF scan with optimal conditions of Δλ = 60 nm. We wished to utilize SF to explore whether lysed cells and diffused molecules from the substrate limit the detection of *φx*-*174* based on its Trp content. *φx*-*174* plaques were replicated on an E. *coli* layer on top of NA agar. The *φx*-*174* were extracted by ammonium acetate buffer for 10 min. The supernatant was removed and filtrated through an 0.22 μm filter and again by 0.02 μm. The same procedure was repeated for Petri dishes without *φx-174.* To evaluate the Trp concentration of the filtrates, SF measurements were performed on the Alumina filter (Δλ = 60 nm, λ_em_ (280 – 450 nm) (Fig. 1). The ratio between the maximal intensities of the filtrates (λ_Em_ = 345 nm) without and with *φx*-*174* was 68.8%. These results shows that the high fluorescence contribution of the NA or lysed bacterial cells highly overlaps with the fluorescence that originates from *φx-174.* This overlap limits the ability to make a reliable detection based the Trp content of the virus.

**Fig. 1.**
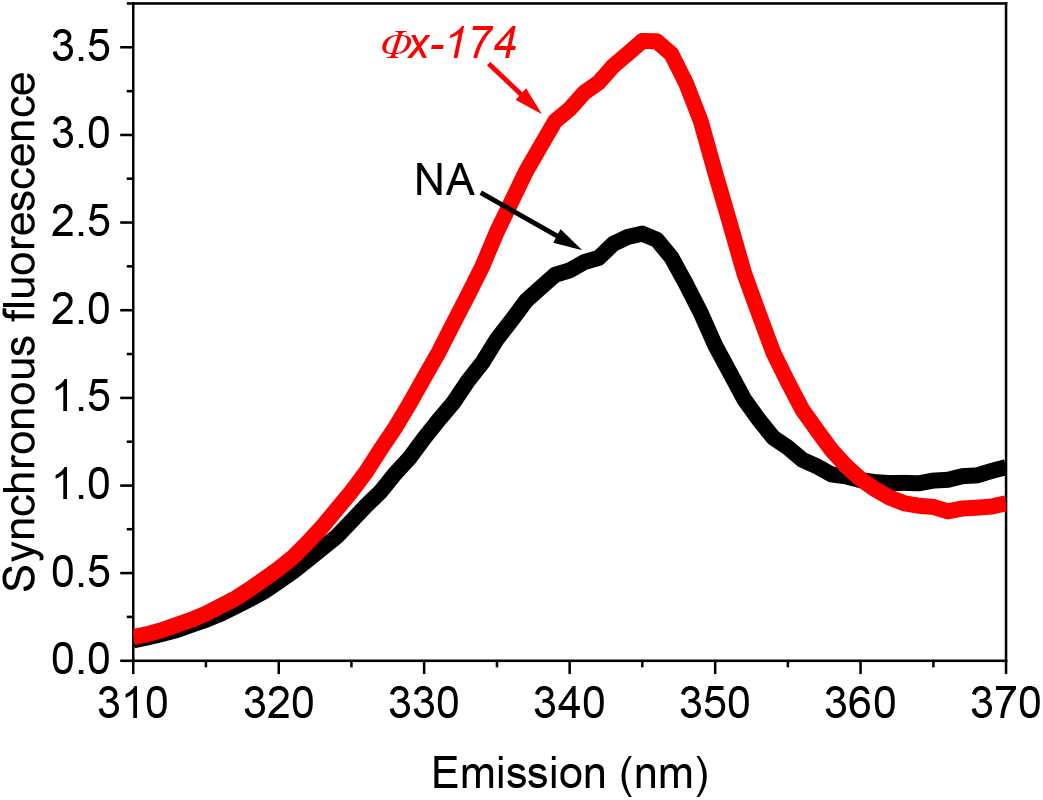
Tryptophan residues in the bacterial media limits the detection of *φx*-*174*. SF spectra of E. *coli* extracts on Alumina filter after filtration with and without *φx*-*174* (Δλ = 60 nm, λ_Em_ = 310 – 370 nm). SF of extracts without *φx*-*174* (black), and with *φx*-*174* (red).

### Preparation of a cultivation medium with low amino acids content

To decrease the Trp fluorescence contribution that derives from the bacterial cultivation medium, we prepared a new medium which consist of less Trp than in NA. The main ingredients of the low Trp medium (LTM) were Agar, glucose, a salts mixture and yeast extract (for more details see experimental section). E. *coli* was successfully cultivated on LTM to form a uniform layer.

Next, we wished to estimate the spectral interference of each individual group of ingredients and the mixture of all ingredients on spectral overlap with Trp. Media with Agar, Agar + Glucose, Agar + salts, Agar + Yeast extracts and Agar + Glucose + Salts + Yeast extracts (YE) were prepared. Ammonium acetate buffer was added on the top of the substrates and incubated for 10 min. The buffer was removed and filtrated. SF spectra of the filtrates (Δλ = 60 nm, λ_Em_ = 310 – 370 nm) were measured (Fig. 2). The filtrates of the Agar and Agar + Glucose extraction did not have a significant fluorescence contribution. A small fluorescence contribution of 0.1 CPS was obtained from the Agar + Salts extract. However, the shape of the peak was not similar to the fluorescence peak of Trp. A peak with a significant intensity of 0.6 CPS was obtained for the YE extract. The intensity of this peak is ~ 5 times less than that obtained when using NA media (Fig. 1). Interestingly, when all ingredients were added to the substrate, no significant fluorescence contribution was observed at the emission wavelengths range of Trp (310 – 370 nm). We postulated that the presence of sugars and salts increases the viscosity of the substrate, and in this way, significantly decreases the diffusion rate of Trp (and other components) from the cultivation media to the ammonium acetate extraction buffer.

**Fig. 2.**
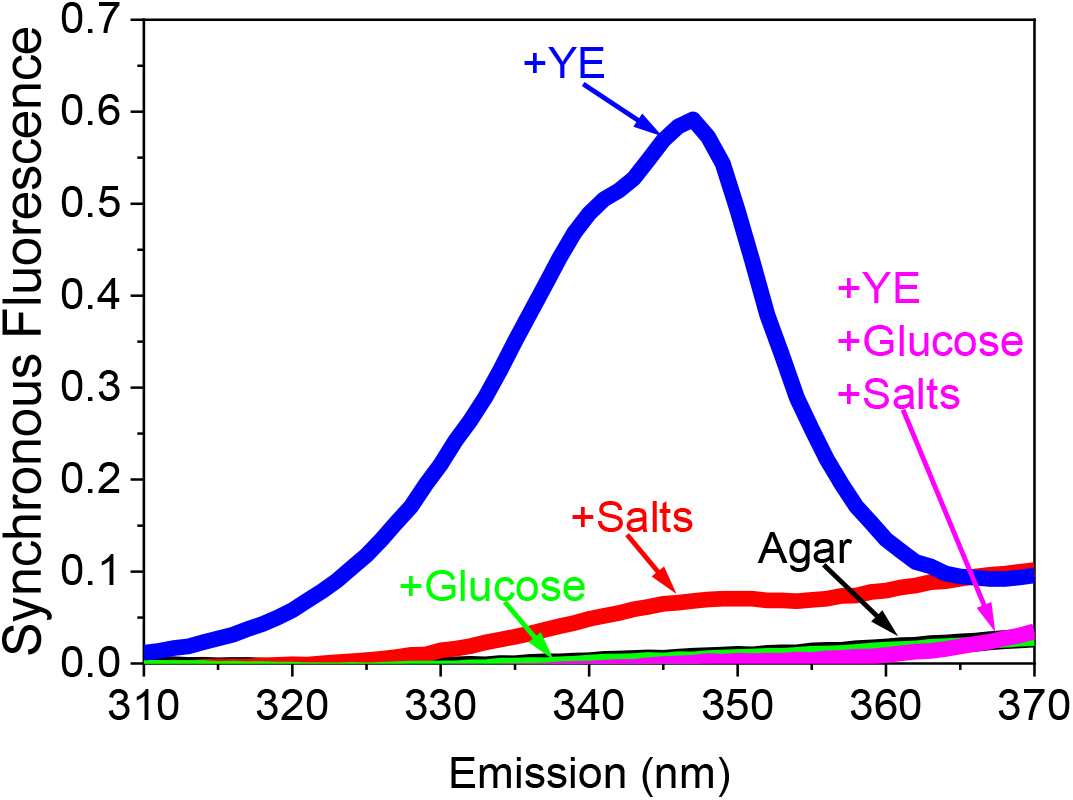
SF measurements of the extracts of the ingredients of the LTM medium. SF of the ingredients of LTM were measured in order to evaluate their fluorescence overlap with TRP. Agar (black), Agar + salts (red), Agar + glucose (green), Agar + YE (blue), Agar + glucose + yeast extract (purple).

### Detection of φx-174 based on its Trp fluorescence fingerprint

Using the LTM reduced the extraction of proteins from the substrate to the ammonium acetate buffer. However, proteins and other fluorescent materials may be secreted from the E. *coli* cells or be released to the buffer by dead or lysed cells. We wished to assess whether such materials are extracted into the Ammonium acetate buffer and whether their fluorescence overlaps with the spectral fingerprint of Trp. For this reason, E. *coli* was cultivated on the LTM with or without the addition of *φx-174.* Ammonium acetate buffer was added on the top of the LTM for 10 min. Then, the buffer was removed and filtrated through 0.22 μm filter, and SF of the filtrates was measured (Fig. 3). A fluorescence contribution between 330 – 370 nm was obtained from the sample without *φx*-*174*. However, its spectral shape was linear and clearly did not originate from Trp, while the samples with *φx*-*174* showed the spectral fingerprint of Trp. We suggest that this fluorescence contribution derives from residues of NAD^+^, and NADH molecules that may overlap if the concentrations are high enough ^46^. We suggest that the appearance of these molecules derive cells that were lysed by *φx*-*174* spilling their NADH pools into the external media. The obtained result show that when using the LTM substrate (instead of the conventional NA), it is possible to use SF to detect the Trp fluorescence contribution which originates only from φx-174 and not from the host cultivation medium.

**Fig. 3.**
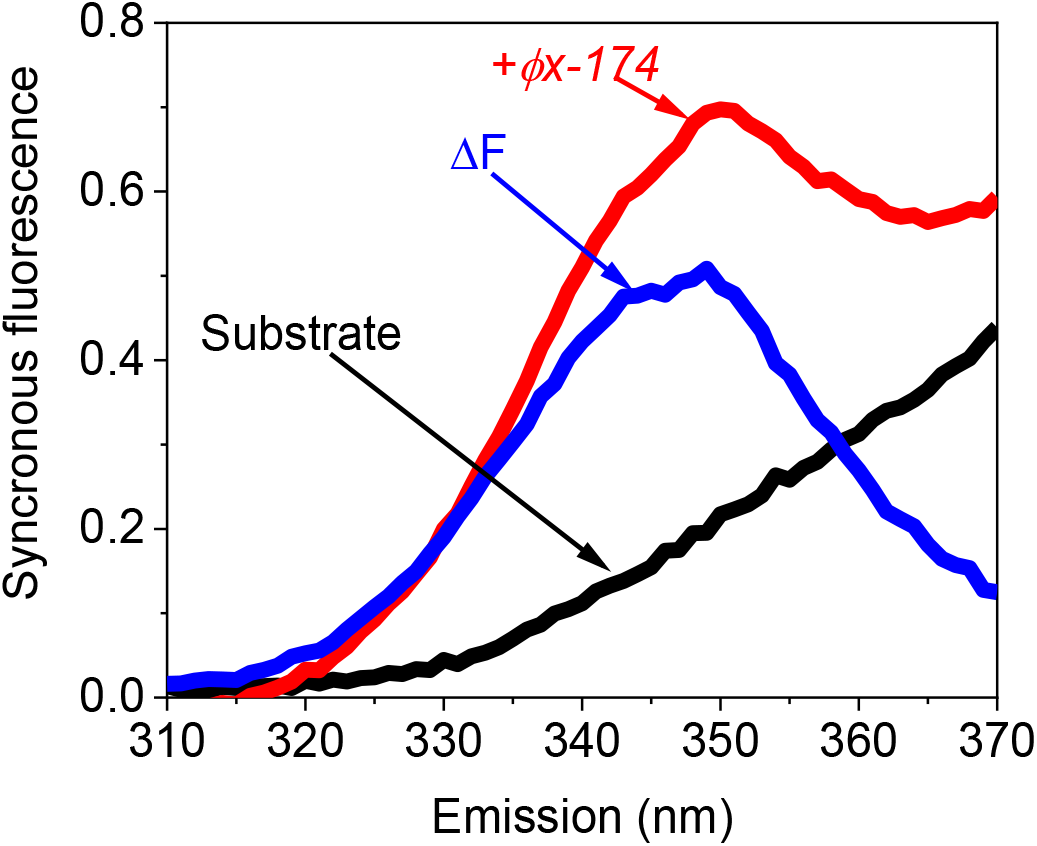
Detection of φx-174 based on its Trp fluorescence fingerprint. SF spectra of the extraction of LTM into the ammonium acetate buffer, without (black), without (red) *φx*-*174*, and their mathematical subtraction (ΔF, blue).

To evaluate the Trp concentration, the SF spectra of the sample without *φx*-*174* was subtracted from the sample with it. Calibration curves of concentration versus the intensity at λ_em_ = 350 nm were prepared by measuring SF spectra of Trp. The concentration of Trp in *φx*-*174* was determined to be 4.4 −/+ 0.3 nM.

A plaque assay was conducted to evaluate the number of virus copies in the extraction buffer (for more detailed explanation, see materials & methods). Based on the plaque assay, the concentration of Trp per virus was evaluated to be 4.4 zeptomol / PFU (that is equivalent to 2650 Trp units).

To compare this experimental result with the theoretical number of Trp copies, we performed a bioinformatic analysis based on the deposited protein sequences that are deposited in the protein data bank (Fig. S1). In this analysis, we summarized the total number of Trp units in all of the copies of the main proteins in *φx-174:* capsid protein F, major spike protein G, minor spike protein H, and DNA-binding protein J. The calculated number of Trp units was 456 was about 5 times lesser than obtained in our experiments. We suggest that this difference may derive from the Forster Energy Transfer (FET) between Tyrosine and Trp that are closely associated in the virus particle. This FET may enhance the fluorescence intensity of Trp ^47^

### A model for the method of the *φx*-*174* purification and detection

Based on the presented results, we propose a mechanism that explains the basic principles of the replication, purification, and detection of *φx*-*174* based on the utilization of LTM and the application of SF.

## Conclusions

In this work, we present a new method for fast screening of viral particles grown on bacterial substrates. We developed a formulation for bacterial cultivation medium which lowers the concentration of Trp in comparison to NA, which is the frequently used cultivation medium. We show that by using this substrate as a platform for cultivation of *E. coli* and replication of the virus φx-174, it is possible to detect *φx*-*174* based on its intrinsic fluorescence which originates from its Trp content. This separation and detection method is simple and rapid and may be used in the future for the identification of various viral species. We demonstrate for the first time the possibility of using intrinsic fluorescence to detect a virus. The concept of coupling intrinsic fluorescence-based methods to decreasing the impurities in the source, may be implemented for rapid detecting of pathogenic virus.

## Acknowledgements

This study has been supported by an internal fund (Micro Grants for the Technion Recycling Initiative, Technion, Israel). Yaniv Shlosberg is supported by the Schulich Graduate fellowship. We thank Prof. Noam Adir for his scientific consultation in the writing of the paper.

## Author contributions

YF and YS conceived the idea. YF and YS designed the experiments. YF and YS performed the main experiments. YF and YS wrote the paper. IS and RA supervised the entire research project and provide funding.

## References

(1) Salemi, S.; Markovic, M.; Martini, G.; D’Amelio, R. The Expanding Role of Therapeutic Antibodies. Int. Rev. Immunol. 2015, 34 (3), 202–264. https://doi.org/10.3109/08830185.2013.863304.

(2) Li, C.; Mori, L.; Valente, S. T. The Block-and-Lock Strategy for Human Immunodeficiency Virus Cure: Lessons Learned from Didehydro–Cortistatin A. J. Infect. Dis. 2021, 223 (Supplement_1), S46–S53. https://doi.org/10.1093/infdis/jiaa681.

(3) Komiyama, M. Molecular-Level Anatomy of SARS-CoV-2 for the Battle against the COVID-19 Pandemic. Bull. Chem. Soc. Jpn. 2021, 94 (5), 1478–1490. https://doi.org/10.1246/bcsj.20210030.

(4) Arias-Arias, J. L.; MacPherson, D. J.; Hill, M. E.; Hardy, J. A.; Mora-Rodríguez, R. A Fluorescence-Activatable Reporter of Flavivirus NS2B–NS3 Protease Activity Enables Live Imaging of Infection in Single Cells and Viral Plaques. J. Biol. Chem. 2020, 295 (8), 2212–2226. https://doi.org/10.1074/jbc.RA119.011319.

(5) Crosson, S. M.; Dib, P.; Smith, J. K.; Zolotukhin, S. Helper-Free Production of Laboratory Grade AAV and Purification by Iodixanol Density Gradient Centrifugation. Mol. Ther. - Methods Clin. Dev. 2018, 10, 1–7. https://doi.org/10.1016/j.omtm.2018.05.001.

(6) Hadiji-Abbes, N.; Martin, M.; Benzina, W.; Karray-Hakim, H.; Gergely, C.; Gargouri, A.; Mokdad-Gargouri, R. Extraction and Purification of Hepatitis B Virus-like M Particles from a Recombinant Saccharomyces Cerevisiae Strain Using Alumina Powder. J. Virol. Methods 2013, 187 (1), 132–137. https://doi.org/10.1016/j.jviromet.2012.09.023.

(7) Rocha, J. M. Aqueous Two-Phase Systems and Monolithic Chromatography as Alternative Technological Platforms for Virus and Virus-like Particle Purification. J. Chem. Technol. \& Biotechnol. 2021, 96 (2), 309–317. https://doi.org/https://doi.org/10.1002/jctb.6595.

(8) Steger, G.; Riesner, D. Viroid Research and Its Significance for RNA Technology and Basic Biochemistry. Nucleic Acids Res. 2018, 46 (20), 10563–10576. https://doi.org/10.1093/nar/gky903.

(9) Popov, V. L.; Tesh, R. B.; Weaver, S. C.; Vasilakis, N. Electron Microscopy in Discovery of Novel and Emerging Viruses from the Collection of the World Reference Center for Emerging Viruses and Arboviruses (WRCEVA). Viruses 2019, 11 (5). https://doi.org/10.3390/v11050477.

(10) Mukherjee, S.; Boutant, E.; Réal, E.; Mély, Y.; Anton, H. Imaging Viral Infection by Fluorescence Microscopy: Focus on HIV-1 Early Stage. Viruses 2021, 13 (2). https://doi.org/10.3390/v13020213.

(11) Carter and Saunders. Virology: Principles and Applications (2nd Ed.); 2012.

(12) Gensberger, E. T.; Kostić, T. Novel Tools for Environmental Virology. Current Opinion in Virology. Elsevier B.V. February 2013, pp 61–68. https://doi.org/10.1016/j.coviro.2012.11.005.

(13) Cheng, M. P.; Papenburg, J.; Desjardins, M.; Kanjilal, S.; Quach, C.; Libman, M.; Dittrich, S.; Yansouni, C. P. Diagnostic Testing for Severe Acute Respiratory Syndrome-Related Coronavirus 2: A Narrative Review. Annals of internal medicine. NLM (Medline) June 2020, pp 726–734. https://doi.org/10.7326/M20-1301.

(14) Ménard-Moyon, C.; Bianco, A.; Kalantar-Zadeh, K. Two-Dimensional Material-Based Biosensors for Virus Detection. ACS Sensors 2020, 5 (12), 3739–3769. https://doi.org/10.1021/acssensors.0c01961.

(15) Banerjee, S.; Maurya, S.; Roy, R. Single-Molecule Fluorescence Imaging: Generating Insights into Molecular Interactions in Virology. J. Biosci. 2018, 43 (3), 519–540. https://doi.org/10.1007/s12038-018-9769-y.

(16) Chojnacki, J.; Eggeling, C. Super-Resolution Fluorescence Microscopy Studies of Human Immunodeficiency Virus. Retrovirology. BioMed Central Ltd. June 2018, p 41. https://doi.org/10.1186/s12977-018-0424-3.

(17) De Almeida Pondé, R. A. Enzyme-Linked Immunosorbent/Chemiluminescence Assays, Recombinant Immunoblot Assays and Nucleic Acid Tests in the Diagnosis of HCV Infection. European Journal of Clinical Microbiology and Infectious Diseases. August 2013, pp 985–988. https://doi.org/10.1007/s10096-013-1857-1.

(18) Iha, K.; Inada, M.; Kawada, N.; Nakaishi, K.; Watabe, S.; Tan, Y. H.; Shen, C.; Ke, L.-Y.; Yoshimura, T.; Ito, E. Ultrasensitive ELISA Developed for Diagnosis. Diagnostics 2019, 9 (3), 78. https://doi.org/10.3390/diagnostics9030078.

(19) Cantalupo, P. G.; Pipas, J. M. Detecting Viral Sequences in NGS Data. Curr. Opin. Virol. 2019, 39, 41–48. https://doi.org/https://doi.org/10.1016/j.coviro.2019.07.010.

(20) Tisthammer, K. H.; Dong, W.; Joy, J. B.; Pennings, P. S. Comparative Analysis of Within-Host Mutation Patterns and Diversity of Hepatitis Cvirus Subtypes 1a, 1b, and 3a. Viruses 2021, 13 (3). https://doi.org/10.3390/v13030511.

(21) Srivastava, S.; Upadhyay, D. J.; Srivastava, A. Next-Generation Molecular Diagnostics Development by CRISPR/Cas Tool: Rapid Detection and Surveillance of Viral Disease Outbreaks. Front. Mol. Biosci. 2020, 7, 582499. https://doi.org/10.3389/fmolb.2020.582499.

(22) Singh, S.; Kumar, V.; Kapoor, D.; Dhanjal, D. S.; Bhatia, D.; Jan, S.; Singh, N.; Romero, R.; Ramamurthy, P. C.; Singh, J. Detection and Disinfection of COVID-19 Virus in Wastewater. Environmental Chemistry Letters. Springer Science and Business Media Deutschland GmbH February 2021, p 3. https://doi.org/10.1007/s10311-021-01202-1.

(23) Qian, J.; Boswell, S. A.; Chidley, C.; Lu, Z. xiang; Pettit, M. E.; Gaudio, B. L.; Fajnzylber, J. M.; Ingram, R. T.; Ward, R. H.; Li, J. Z.; Springer, M. An Enhanced Isothermal Amplification Assay for Viral Detection. Nat. Commun. 2020, 11 (1), 1–10. https://doi.org/10.1038/s41467-020-19258-y.

(24) Naveen, K. P.; Bhat, A. I. Development of Reverse Transcription Loop-Mediated Isothermal Amplification (RT-LAMP) and Reverse Transcription Recombinase Polymerase Amplification (RT-RPA) Assays for the Detection of Two Novel Viruses Infecting Ginger. J. Virol. Methods 2020, 282, 113884. https://doi.org/10.1016/j.jviromet.2020.113884.

(25) Obande, G. A.; Singh, K. K. B. Current and Future Perspectives on Isothermal Nucleic Acid Amplification Technologies for Diagnosing Infections. Infection and Drug Resistance. Dove Medical Press Ltd. 2020, pp 455–483. https://doi.org/10.2147/IDR.S217571.

(26) Shahzad, A.; Edetsberger, M.; Koehler, G. Fluorescence Spectroscopy: An Emerging Excellent Diagnostic Tool in Medical Sciences. Appl. Spectrosc. Rev. 2010, 45 (1), 1–11. https://doi.org/10.1080/05704920903435375.

(27) Ruokola, P.; Dadu, E.; Kazmertsuk, A.; Hakkanen, H.; Marjomaki, V.; Ihalainen, J. A. Raman Spectroscopic Signatures of Echovirus 1 Uncoating. J. Virol. 2014, 88 (15), 8504–8513. https://doi.org/10.1128/jvi.03398-13.

(28) Lambert, P. J.; Whitman, A. G.; Dyson, O. F.; Akula, S. M. Raman Spectroscopy: The Gateway into Tomorrow’s Virology. Virology Journal. BioMed Central June 2006, p 51. https://doi.org/10.1186/1743-422X-3-51.

(29) Santos, M. C. D.; Morais, C. L. M.; Nascimento, Y. M.; Araujo, J. M. G.; Lima, K. M. G. Spectroscopy with Computational Analysis in Virological Studies: A Decade (2006–2016). TrAC - Trends in Analytical Chemistry. Elsevier B.V. December 2017, pp 244–256. https://doi.org/10.1016/j.trac.2017.09.015.

(30) Shen, F.; Triezenberg, S. J.; Hensley, P.; Porter, D.; Knutson, J. R. Critical Amino Acids in the Transcriptional Activation Domain of the Herpesvirus Protein VP16 Are Solvent-Exposed in Highly Mobile Protein Segments: An Intrinsic Florescence Study. J. Biol. Chem. 1996, 271 (9), 4819–4826. https://doi.org/10.1074/jbc.271.9.4819.

(31) Lakowicz, J. R. Fluorophores. In Principles of Fluorescence Spectroscopy; Springer US, 2006; pp 63–95. https://doi.org/10.1007/978-0-387-46312-4_3.

(32) Marinowic, D. R.; Zanirati, G.; Rodrigues, F. V. F.; Grahl, M. V. C.; Alcará, A. M.; Machado, D. C.; Da Costa, J. C. A New SYBR Green Real-Time PCR to Detect SARS-CoV-2. Sci. Rep. 2021, 11 (1), 2224. https://doi.org/10.1038/s41598-021-81245-0.

(33) Andrade-Eiroa, Á.; de-Armas, G.; Estela, J. M.; Cerdà, V. Critical Approach to Synchronous Spectrofluorimetry. I. TrAC - Trends in Analytical Chemistry. Elsevier September 2010, pp 885–901. https://doi.org/10.1016/j.trac.2010.04.010.

(34) Liu, Q.; Grant, G.; Vo-Dinh, T. Investigation of Synchronous Fluorescence Method in Multicomponent Analysis in Tissue. IEEE J. Sel. Top. Quantum Electron. 2010, 16 (4), 927–940. https://doi.org/10.1109/JSTQE.2009.2031162.

(35) Rubio, S.; Gomez-Hens, A.; Valcarcel, M. Analytical Applications of Synchronous Fluorescence Spectroscopy. Talanta 1986, 33 (8), 633–640. https://doi.org/https://doi.org/10.1016/0039-9140(86)80149-7.

(36) Lloyd, J. B. F.; Evett, I. W. Prediction of Peak Wavelengths and Intensities in Synchronously Excited Fluorescence Emission Spectra. Anal. Chem. 1977, 49 (12), 1710–1715. https://doi.org/10.1021/ac50020a020.

(37) Patra, D.; Mishra, A. K. Recent Developments in Multi-Component Synchronous Fluorescence Scan Analysis. TrAC Trends Anal. Chem. 2002, 21 (12), 787–798. https://doi.org/https://doi.org/10.1016/S0165-9936(02)01201-3.

(38) Lloyd, J. B. F. Synchronized Excitation of Fluorescence Emission Spectra. Nat. Phys. Sci. 1971, 231 (20), 64–65. https://doi.org/10.1038/physci231064a0.

(39) Ammor, M. S. Recent Advances in the Use of Intrinsic Fluorescence for Bacterial Identification and Characterization. J. Fluoresc. 2007, 17 (5), 455–459. https://doi.org/10.1007/s10895-007-0180-6.

(40) Perinchery, S. M.; Kuzhiumparambil, U.; Vemulpad, S.; Goldys, E. M. The Potential of Autofluorescence Spectroscopy to Detect Human Urinary Tract Infection. Talanta 2010, 82 (3), 912–917. https://doi.org/https://doi.org/10.1016/j.talanta.2010.05.049.

(41) Sahar, A.; Boubellouta, T.; Dufour, É. Synchronous Front-Face Fluorescence Spectroscopy as a Promising Tool for the Rapid Determination of Spoilage Bacteria on Chicken Breast Fillet. Food Res. Int. 2011, 44 (1), 471–480. https://doi.org/https://doi.org/10.1016/j.foodres.2010.09.006.

(42) Sohn, M.; Himmelsbach, D. S.; Barton, F. E.; Fedorka-Cray, P. J. Fluorescence Spectroscopy for Rapid Detection and Classification of Bacterial Pathogens. Appl. Spectrosc. 2009, 63 (11), 1251–1255. https://doi.org/10.1366/000370209789806993.

(43) Li, R.; Goswami, U.; King, M.; Chen, J.; Cesario, T. C.; Rentzepis, P. M. In Situ Detection of Live-to-Dead Bacteria Ratio after Inactivation by Means of Synchronous Fluorescence and PCA. Proc. Natl. Acad. Sci. U. S. A. 2018, 115 (4), 668–673. https://doi.org/10.1073/pnas.1716514115.

(44) Kropinski, A. M.; Mazzocco, A.; Waddell, T. E.; Lingohr, E.; Johnson, R. P. Enumeration of Bacteriophages by Double Agar Overlay Plaque Assay. Methods Mol. Biol. 2009, 501, 69–76. https://doi.org/10.1007/978-1-60327-164-6_7.

(45) Shlosberg, Y.; Farber, Y.; Hasson, S.; Bulatov, V.; Schechter, I. Fast Label-Free Identification of Bacteria by Synchronous Fluorescence of Amino Acids. Anal. Bioanal. Chem. 2021. https://doi.org/10.1007/s00216-021-03642-8.

(46) Shlosberg, Y.; Eichenbaum, B.; Tóth, T. N.; Levin, G.; Liveanu, V.; Schuster, G.; Adir, N. NADPH Performs Mediated Electron Transfer in Cyanobacterial-Driven Bio-Photoelectrochemical Cells. iScience 2020, 24 (1), 101892.

(47) zhang, Y.; Yang, X.; Liu, L.; Huang, X.; Pu, J.; Long, G.; Zhang, L.; Liu, D.; Xu, B.; Liao, J.; Liao, F. Comparison of Förster-Resonance-Energy-Transfer Acceptors for Tryptophan and Tyrosine Residues in Native Proteins as Donors. J. Fluoresc. 2013, 23 (1), 147–157. https://doi.org/10.1007/s10895-012-1128-z.

